# Metaproteomics of colonic microbiota unveils discrete protein functions among colitic mice and control groups

**DOI:** 10.1101/219782

**Authors:** Clara Moon, Gregory S. Stupp, Andrew I. Su, Dennis W. Wolan

## Abstract

Metaproteomics can greatly assist established high-throughput sequencing methodologies to provide systems biological insights into the alterations of microbial protein functionalities correlated with disease-associated dysbiosis of the intestinal microbiota. Here, we utilized the well-characterized murine T cell transfer model of colitis to find specific changes within the intestinal luminal proteome associated with inflammation. MS proteomic analysis of colonic samples permitted the identification of ∽10,000-12,000 unique peptides that corresponded to 5,610 protein clusters identified across three groups, including the colitic *Rag1*^-/-^ T cell recipients, isogenic *Rag1*^-/-^ controls, and wild-type mice. We demonstrate that the colitic mice exhibited a significant increase in *Proteobacteria* and *Verrucomicrobia* and show that such alterations in the microbial communities contributed to the enrichment of specific proteins with transcription and translation gene ontology terms. In combination with 16S sequencing, our metaproteomics-based microbiome studies provide a foundation for assessing alterations in intestinal luminal protein functionalities in a robust and well-characterized mouse model of colitis, and set the stage for future studies to further explore the functional mechanisms of altered protein functionalities associated with dysbiosis and inflammation.

**Statement of significance of the study:** The commensal gut microbiota is essential to maintaining health and has a primary role in digestion/metabolism, homeostasis, and protection from pathogenic bacteria. High-throughput sequencing has established *Bacteroidetes, Firmicutes, Proteobacteria,* and *Actinobacteria* as the four major bacterial phyla that comprise the ecological makeup of the intestinal microbiota. However, the tremendous inter-/intra-variability in microbial composition across individuals, as well as along the length of the intestinal tract has made it difficult to definitively ascertain specific bacterial species associated with health or as drivers of disease states, including inflammatory bowel disease. In this study, we expanded upon the current metaproteomics techniques and use the robust and highly reproducible murine T cell transfer model of colitis as well as a comprehensive database of mouse, human, plant, and all microbial genomes sequenced to date to elucidate alterations in both host and gut microbial proteins associated with intestinal inflammation. Our results show that host genetics, gut microbiota, and inflammation have tremendous influences on the intestinal luminal proteomic landscape.

## 1 Introduction

IBD is a complex disorder caused by many variables, including host genetics, environmental factors, and the intestinal microbiota [1-6]. Numerous mouse models of colitis suggest an important role for the intestinal bacteria in disease propagation. For example, treatment of colitic mice with antibiotics or raising such inflammation-prone mice in a germ-free environment result in the amelioration and/or prevention of disease [7,8]. These results readily translate into the clinic, as human IBD patients are frequently prescribed antibiotics, such as metronidazole (Flagyl) and ciprofloxacin [9]. Accordingly, alterations in the composition of the normal commensal gut microbiota, known as dysbiosis, is a major contributing factor to disease [10,11]. Metagenomics-based approaches have demonstrated that decreases in the *Bacteroidetes* and *Firmicutes* phyla, and increases in *Proteobacteria* and *Actinobacteria* are globally altered in IBD [12]. However, specific species with biological relevance to driving the disease phenotype have yet to be identified due to the inter-/intra-individual variation in microbial diversity.

We and others have developed technological advances in metaproteomics to interrogate and quantitate differences in microbial proteins, as protein functions are well-conserved across commensal organisms and their hosts despite the tremendous ecological diversity within gut microbiomes [13,14]. Such key studies employing mass spectrometry have laid the foundation for the field of microbiome metaproteomics [15,16]. Recently, comparisons between healthy and IBD human patient samples have been published. For example, Erickson *et al.* compared fecal samples from 6 twin pairs that included healthy, ileal Crohn’s Disease (CD), and colonic CD patients. Metaproteomic data collected by liquid chromatography coupled with tandem mass spectrometry (LC-MS/MS) identified 700-1,250 or 1,900-3,000 protein clusters within the fecal samples when searched against the individual’s matched metagenome or 51 human microbial isolate reference genomes, respectively [17,18]. The integration of metagenomics sequencing and reference genome sequences of common gut bacteria to assess metaproteomic data permitted this study to identify significant alterations in protein functionalities differentially represented in the healthy vs. ileal CD patients *((e.g.,* proteogenomics).

In this study, we expand upon the current metaproteomics techniques and employed <u>mul</u>ti<u>d</u>imensional protein identification technology (MudPIT) [19] and the recently described <u>Com</u>prehensive <u>P</u>rotein <u>I</u>dentification <u>L</u>ibrary (ComPIL) [20] to generate a compendium of host and microbial protein functions that are significantly correlated with intestinal inflammation. Using these methods, we found 10,000-12,000 unique peptides per treatment group that corresponded to a total of 5,610 protein clusters. Bioinformatic analyses of colonic contents isolated from colitic mice and comparison of the metaproteomic composition to two different control groups delineated many unique host and microbial functionalities among all three groups, as well as interesting commonalities. Our results demonstrate and support that a combination of variables, including host genetics, the intestinal gut microbiota, and inflammation importantly contribute to the alterations observed in the intestinal luminal proteome.

## 2 Materials and methods

### 2.1 Mice

Animal protocols were approved by The Institutional Animal Care and Use Committee (IACUC) at The Scripps Research Institute (TSRI); IACUC protocol #:16-0023. 6-week old B6.129S7-*Rag1*^*tm*1*mom*^/*J* (*Rag1*^-/-^) mice (Rag) and wild-type C57BL/6J (WT) mice were cohoused for two weeks to normalize the microbiota (see Supporting Information 1 for details). Briefly, CD3+CD4+CD8^-^CD25^-^Foxp3^-^ naïve T cells from the spleens of donor *Foxp3-EGFP* reporter mice were transferred retro-orbitally to five Rag mice to generate the T cell recipient mice group (“RT” mice), as previously described [21,22]. Control Rag (n=2) and WT (n=5) mice were injected retro-orbitally with a similar volume of sterile PBS (Supporting Information Table 1) and all mice were separated by treatment group for the remainder of the experiment. Mice were euthanized on day 56 post T cell transfer (one mouse sacrificed prior to day 56 upon reaching 80% of initial weight). Colonic contents were collected by flushing with 5 ml of sterile PBS. Samples were filtered, aliquoted, centrifuged at 10,000xg for 10 min, and aspirated. The resulting bacterial pellets were frozen at -80 °C until needed. Segments of small intestinal and colonic tissues were fixed in 10% formalin for histology and crypt heights and the number of crypts per field were measured in a blinded manner (Supporting Information Fig. 1).

**Fig. 1.**
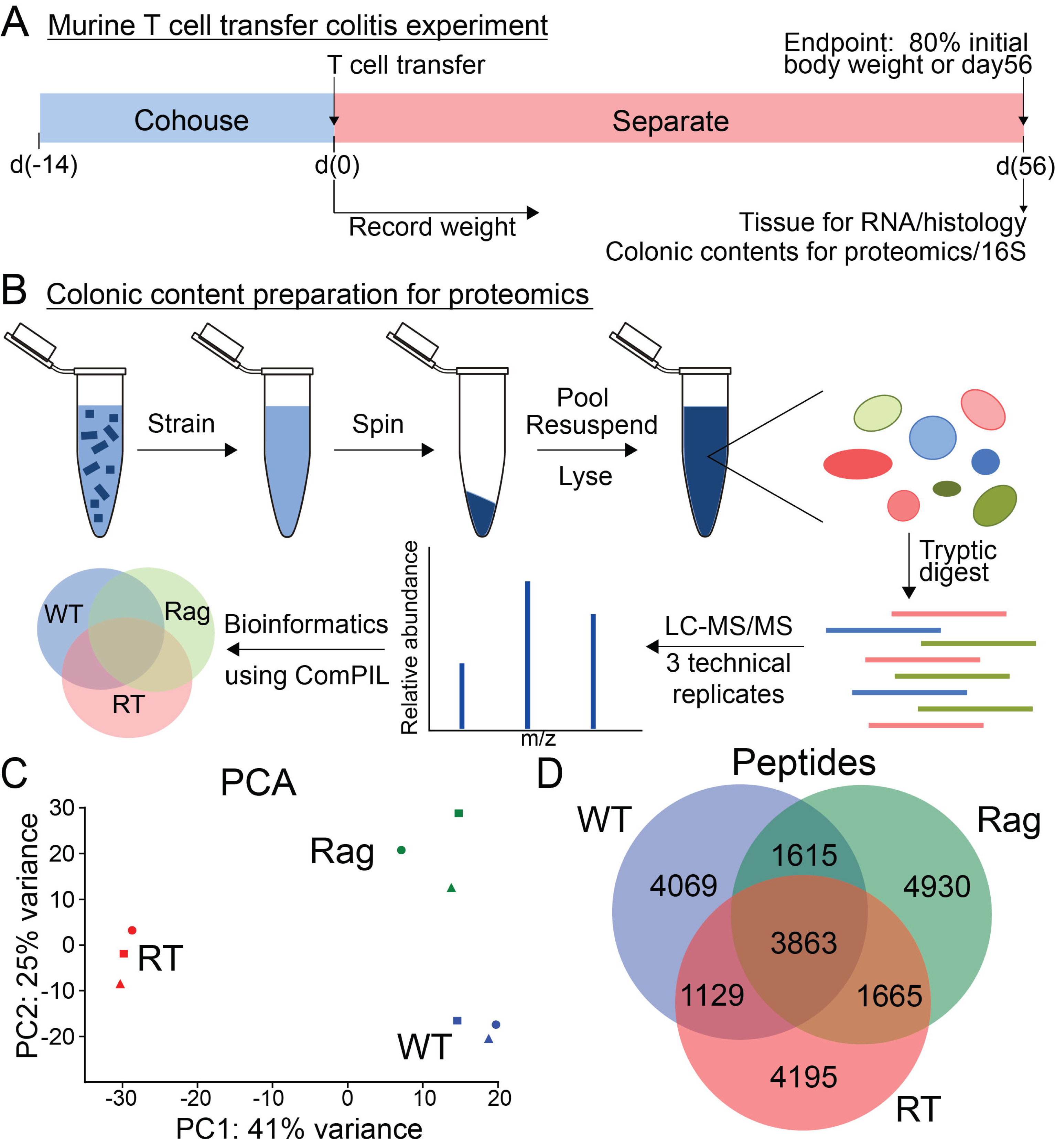
Global proteomic survey of intestinal contents from a murine colitis model. (A) Timeline of the T cell transfer colitis experiment. WT and Rag1^-/-^ mice were initially cohoused for 2 weeks prior to injection of PBS or 2x10^5^ CD3+CD4+CD8^-^CD25^-^Foxp3(EGFP)^-^ splenic T cells on day 0. Mice were then separated by treatment group for the duration of the experiment. Mice were euthanized on day 56, at which time intestinal tissue was taken for histology, and colonic contents were saved and prepared for proteomics. (B) Schematic of colonic content preparation for proteomics. Colonic contents were flushed with sterile PBS, aliquoted, and centrifuged to remove the supernatants. The bacterial pellets were resuspended in lysis buffer and sonicated. The resulting bacterial proteome was trypsin digested and subjected to LC-MS/MS for proteomic data analysis by ComPIL. (C) Principal component analysis of LC-MS/MS samples. Different colors correspond to different treatment groups. Different shapes correspond to different technical replicates: circles = replicate 1, triangles = replicate 2, and squares = replicate 3. (D) Venn diagram showing differences in peptides identified in each treatment group.

### 2.2 Sample Preparation for Unenriched Proteomic Analyses

After all RT mice were confirmed by histology to have inflammation in the ileum and colon (Supporting Information Table 1, Supporting Information Fig. 1), colonic content bacterial samples were thawed on ice, pooled by treatment group and were used immediately for proteomic sample preparation: bacterial pellet samples were re-suspended in 500 µL of lysis buffer (Roche cOmplete protease inhibitor tablet in PBS). Samples were lysed via sonication at 4 °C for 10 min, insoluble cellular material was removed via centrifugation (10,000xg for 5 min), and the remaining soluble protein concentration was measured (Pierce BCA Protein Assay Kit). 100 µg aliquots were subjected to trypsin digestion to generate peptides for MudPIT shotgun proteomics analysis, as previously described and subsequently stored at -20 °C until LC-MS/MS analysis (see Supporting Information 1 for details) [19].

### 2.3 LC-MS/MS

Trypsin-digested peptides were loaded onto a biphasic MudPIT column packed with Aqua C18 resin followed by Partisphere strong cation exchange resin. Standard MudPIT tandem mass spectrometry was performed using a Thermo LTQ Orbitrap XL mass spectrometer. Briefly, peptides were eluted at 0.2 ml/min using an 11-step MudPIT program, as previously described (see Supporting Information 1 for details) [19]. Precursor ions were recorded by scanning in the range of m/z 400.00–1,800.00 with the FTMS analyzer and a resolution of 60,000. The top eight peaks were selected for fragmentation using CID with normalized collision energy set to 35.0. Dynamic exclusion was enabled with a repeat count of 1, repeat duration of 30.00 sec, exclusion list size of 100, and exclusion duration set to 60.00 sec.

### 2.4 Peptide Identification using ComPIL

Precursor and fragmentation ion data were extracted from the Xcalibur RAW files via rawXtract 1.9.9.2 (http://fields.scripps.edu/yates/wp/?pageid=17) in the MS1 and MS2 file formats. The MS2 spectra were scored with Blazmass 0.9993 against peptides of the ComPIL database, containing over 80 million proteins from multiple microbial database sources as well as human, mouse, and plant proteins [20]. Both Blazmass and ComPIL source code are open source (https://github.com/sandipchatterjee/blazmasscompil). Settings for peptide scoring included: 1) a variable modification of oxidized methionine (+15.9949 Da), 2) a static modification for alkylated cysteine residues (+57.02146 Da), 3) a precursor mass tolerance of 10 ppm and 50 ppm tolerance for fragmentation ions, and 4) two missed tryptic cleavages. Filtering was performed using DTASelect 2.1.3 (http://fields.scripps.edu/yates/wp/7pageid=17), requiring 2 peptides per protein and a false discovery rate (FDR) of 1% with respect to peptides [23,24]. FDR was assessed using the target-decoy approach with protein sequences reversed and concatenated with their original protein records [20,25]. The following parameters were used for filtering when run from the command line: “--quiet --brief --trypstat --modstat -y 2 -DM 10 --extra --dm --sfp 0.01 -p 2”.

### 2.5 LC-MS/MS Data Analysis

The source code for this analysis is available online (https://github.com/stuppie/CM7_CM1E2d56col_unenr123_rawextract_2017/). Protein clustering, cluster taxonomy, and gene ontology (GO) term annotations were performed as previously described [20,23]. Briefly, protein loci were mapped to protein clusters using a pre-clustered version of ComPIL using a sequence identity threshold of 70% [20]. A protein cluster was annotated with all GO terms associated with any domain for all possible proteins within that cluster, while removing any GO terms that were parents (“is a” or “part of” relationships) of other GO terms in that protein cluster, using annotations generated from InterProScan v5 (version 5.8-49.0) [23]. Protein cluster differential analysis was performed using DESeq2 1.14.1. The DESeq2 statistical analysis package allows for testing for differential expression in count data, and provides methods designed for dealing with overdispersion, features with low counts, and experiments with low numbers of biological replicates [26]. The method is briefly described as follows: spectral count data is modeled using the negative binomial distribution, which allows determination of the mean-variance relationship. Variance is estimated using an information sharing approach whereby a single feature’s variance is estimated by taking into account information about variances of other similar features measured in the same experiment. Feature significance calling and ranking is then performed by estimating effect sizes, accounting for the logarithmic fold change (LFC) for a feature between treatment and control, and the noisiness of the LFC estimate. The coefficient of variation (CV) for the variance-stabilized transformed spectral counts was calculated by dividing the standard deviation by the mean of spectral counts within each group of technical replicates and the median of the collective CV values is reported as the CV.

### 2.6 Gene set enrichment analysis

Gene set enrichment analysis was performed using gseapy 0.7.6 (https://github.com/BioNinja/gseapy), a Python implementation of the Broad Institute’s Gene Set Enrichment Analysis (GSEA) algorithm [27]. GO gene sets were generated from all identified protein clusters. Terms were subsequently filtered according to the msigdb guidelines: 1) large sets, defined as those containing more than half the total number of protein clusters identified, were removed; 2) sets with less than five members were removed; 3) child terms with the exact same protein cluster members as their parent were removed; and 4) sibling terms with the exact same protein cluster members as other siblings were removed to generate 1 sibling. Significantly altered protein clusters (at a default p-value of <0.20) were ranked by DESeq2-determined shrunken LFC and analyzed. Significantly altered gene sets were called at an adjusted p-value of <0.05.

### 2.7 Taxonomy analysis from proteomic data

Peptide spectral counts were normalized across all samples by a normalization factor of the total number of counts for one experiment divided by the median across all LC-MS/MS experiments, as previously described [23]. Briefly, the peptide taxonomy search space was restricted to the proteins identifiable in a given sample and analysis was performed at the phylum level. Each peptide was traced back to the phylum and if uniquely classifiable, the peptide was classified with a weight of normalized counts. Peptides without a discernible phylum (*e.g.,* could belong more than one) were discarded from analysis. The normalized counts were then used to determine an approximate fractional taxonomic makeup of the sample.

### 2.8 16S rDNA Illumina sequencing and analysis

Colonic content bacterial samples from mice were collected as described above. Genomic DNA was extracted using the Zymo Research Fecal DNA Mini Prep Kit (#11-322) according to manufacturer’s instructions. Genomic DNA samples were submitted to the Scripps Research Next Generation Sequencing Core Facility for preparation of multiplexed amplicon libraries using the NEXTflex™ 16S V4 Amplicon-Seq Kit 2.0 (#4203-03), and sequencing using the 2x300-base pair protocol with the Illumina MiSeq platform. All analysis was performed using Quantitative Insights Into Microbial Ecology (QIIME, version 1.9.1) [28]. Paired-end reads were assembled and quality screened, and had primers removed using PANDAseq 2.10 [29]. Only sequences with length >200 bp were saved. The reads were clustered into operational taxonomic units (OTUs) using the open reference protocol using Greengenes 13.8 [30]. Alpha and beta diversity analyses were conducted on data rarefied to 10,000 sequences per sample. Samples were clustered using the UPGMA method with weighted UniFrac as the distance metric [31].

### 2.9 Availability of data and materials

The datasets generated and/or analyzed during the current study are available in this published article and its supplementary information files, as well as online. Analysis Code can be found at: https://github.com/stuppie/CM7CM1E2d56colunenr123rawextract2017/. The mass spectrometry proteomics data have been deposited to the ProteomeXchange Consortium (http://proteomecentral.proteomexchange.org) via the PRIDE partner repository [32] with the project accession identifier PXD006384.

## 3 Results

### 3.1 Mouse Model of Colitis

We used the well-established murine T cell transfer model of colitis for our studies [21,22]. For these experiments, splenic T cells lacking the regulatory T cell compartment were transferred into immunodeficient *Rag1*^-/-^ recipient mice. Over the course of 6-8 weeks, the RT mice developed robust inflammation along the length of the intestinal tract. Because these mice were not littermates, we initially cohoused the *Rag1*^-/-^ mice with WT C57BL/6 mice for 2 weeks prior to T cell transfer (or PBS control) to account for any microbiota differences in breeding colonies (Fig. 1a). In addition, we designated WT mice and Rag mice as “healthy” control groups for all our proteomic comparisons (Supporting Information Table 1). Upon T cell injection, we separated the mice by treatment group to permit the disease to progress in the RT mice unabated without the potential transfer of “healthy” microbes from the control groups (and vice versa). At the 8-week endpoint, we collected flushed colonic contents for metaproteomic and 16S sequencing analyses, as well as the host intestinal tissue for histology.

### 3.2 Sample Preparation and LC-MS/MS Data Collection

We isolated and prepared intestinal bacteria from colonic content samples for LC-MS/MS proteomic analysis (Fig. 1b). For each treatment group, bacterial pellets were combined and subsequently lysed by sonication, with 5 mice pooled for the WT and RT groups and 2 mice for the Rag group (Supporting Information Table 1). Our goal for this initial study was to focus our metaproteomics data collection and analysis on technical replicates of pooled cohort samples. LC-MS/MS metaproteomics were performed on 3 technical replicates for each pooled treatment group and were highly reproducible, as demonstrated by a CV of 4.1% using the DESeq2 variance-stabilized transformed spectral counts [33]. The resulting proteomic data was searched against the ComPIL database [20,23] and principal component analysis based on the peptide composition among the LC-MS/MS datasets verified that each replicate for the individual treatment groups strongly clustered together (Fig. 1c). Likewise, a hierarchical clustering dendrogram based on the Jaccard distance calculated from the presence or absence of all peptides in a sample further demonstrated that the technical replicates were significantly similar for each treatment group (Supporting Information Fig. 2). In total, we identified 10,676 unique peptides in the WT replicates, 12,073 unique peptides in the Rag replicates, and 10,852 unique peptides in the RT replicates (Fig. 1d, Supporting Information Fig. 3). 3,863 unique peptides were shared across all three cohorts, while each of the treatment groups consisted of ∽4,000 unique and exclusive group-specific peptides (Fig. 1d). Our proteomic data clearly delineated similarities and differences in the intestinal luminal proteome across the three groups. Importantly, the altered peptides were not exclusively variable among the “healthy” and colitic mice, but also between the WT and Rag controls.

**Fig. 2.**
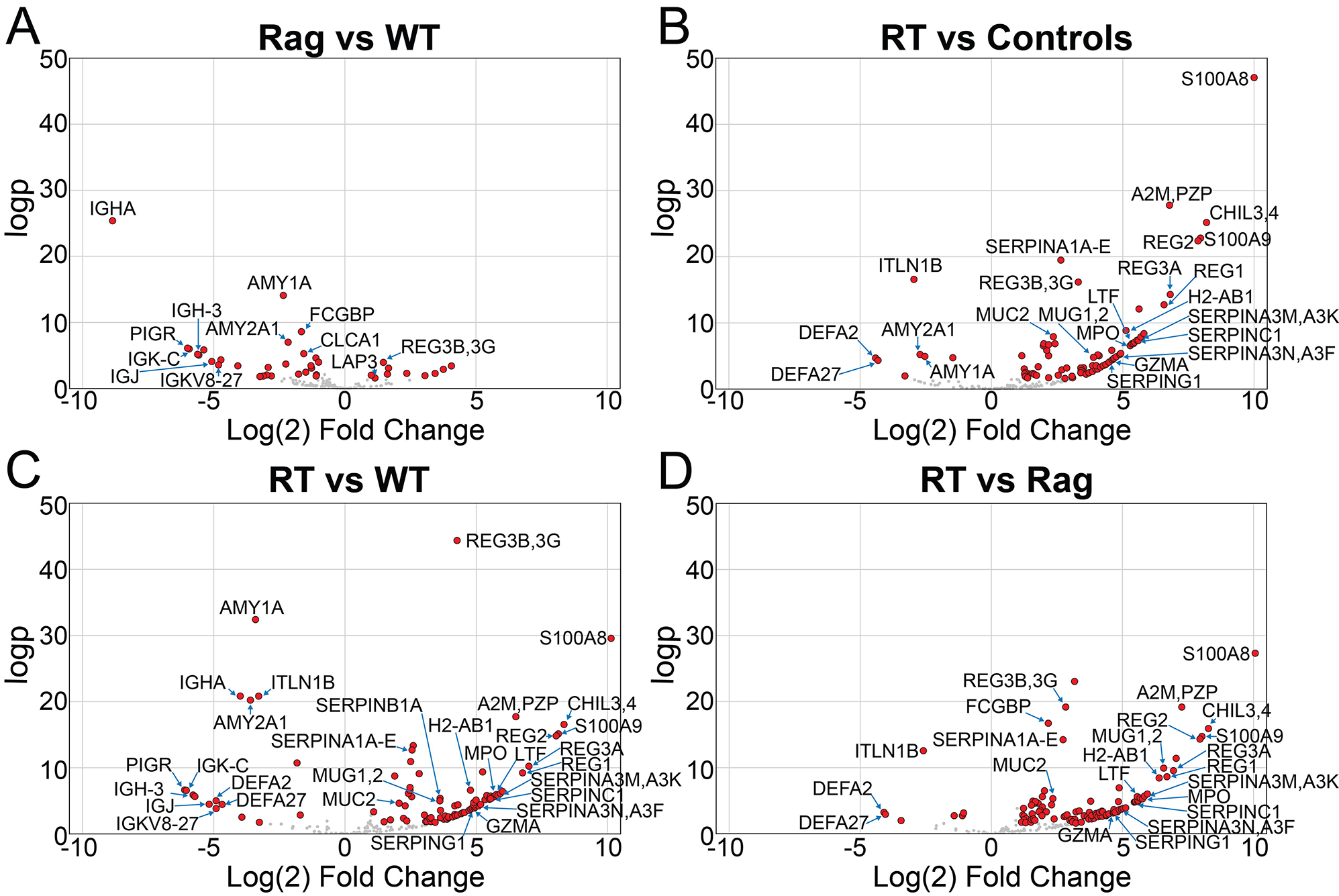
Significantly altered murine protein clusters identified by LC-MS/MS. Volcano plots of murine protein clusters identified in the proteomic samples. The red dots represent protein clusters that are significantly different (FDR <0.05 (logp) and |Log_2_FC| >1) in Rag compared to WT samples (A), RT compared to control samples (B), RT compared to WT samples (C), and RT compared to Rag samples (D). Full lists of the significant differentially expressed proteins and GO terms are available Supporting Information Tables 2, 5, 8, 11.

**Fig. 3.**
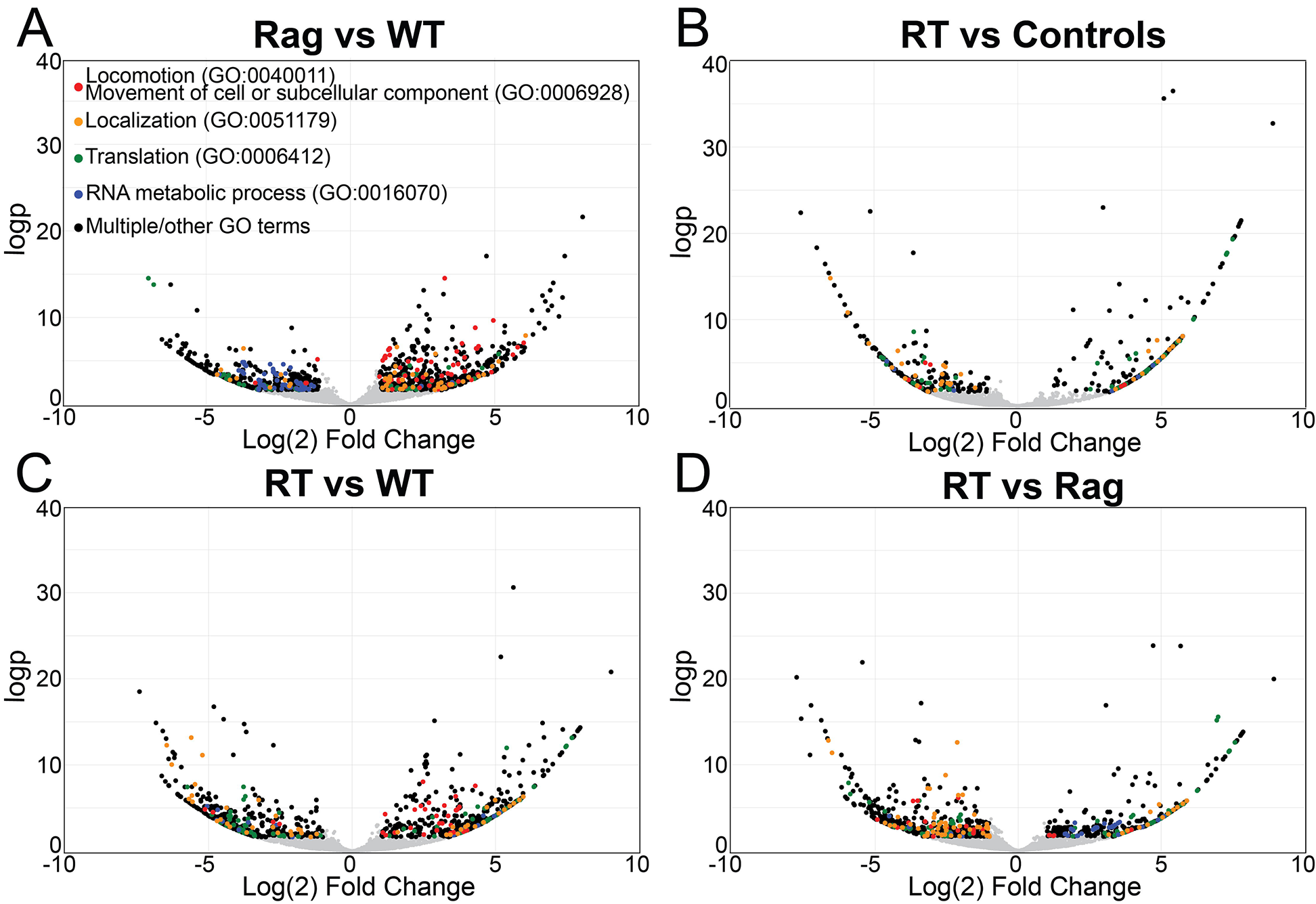
Significantly altered microbial protein clusters identified by LC-MS/MS. Volcano plots of microbial protein clusters identified in the proteomic samples. The colored dots represent the protein clusters that fall under the parent GO terms (and associated child terms) as labeled in the key in A. The black dots represent protein clusters that are significantly different (FDR <0.05 and |Log_2_FC| >1), but do not have assigned GO terms and/or are assigned to multiple/other GO terms. The data compared are as follows: Rag compared to WT samples (A), RT compared to control samples (B), RT compared to WT samples (C), and RT compared to Rag samples (D). Full lists of the significant differentially expressed proteins and GO terms are available Supporting Information Tables 3, 4, 6, 7, 9, 10, 12, 13.

### 3.3 Protein Clusters Differentially Expressed in the Rag and WT Controls

ComPIL was employed to facilitate functional identifications of proteins in each sample group and DESeq2 was used for differential analysis. We first investigated the differences between the Rag and WT control groups and identified a number of protein clusters that were significantly altered (Fig. 2a, Fig. 3a, Supporting Information Tables 2, 3.). Out of the combined 5,610 protein clusters found among all samples, 22 murine and 680 microbial protein clusters were significantly altered with a FDR <0.05 and the absolute value of the log_2_ fold-change (|log_2_FC|) >1. With respect to the altered murine protein clusters, 19 were increased in the WT samples (or missing in Rag mice), while 3 protein clusters were increased in the Rag samples (Fig. 2a, Supporting Information Table 2). Many of the murine proteins identified in these two groups were secreted proteins of either epithelial or immune cell origin. For example, components of the secretory IgA system are increased in the WT samples and absent in the Rag samples, including: 1) the immunoglobulin A heavy chain (IGHA); 2) the J chain (IgJ) which joins the IgA dimers; and 3) the epithelial polymeric immunoglobulin receptor (pIgR) which is cleaved and released along with the dimeric IgA as the secretory component (SC). Additionally, several immunoglobulin light chains as well as the immunoglobulin G heavy chain are also detected only in proteomic samples generated from WT mice. This is expected as the Rag mice lack B cells (and therefore immunoglobulins), thus providing proof-of-principle that MudPIT proteomics of highly complex microbiome proteomes can distinguish differences in host proteins.

The 680 significantly altered microbial protein clusters were equally divided between the WT and Rag groups, with 350 and 330 differential protein clusters, respectively (Fig. 3a, Supporting Information Table 3). GSEA on the microbial protein clusters reveals that the major biological processes (BP) enriched in the Rag mice are locomotion (GO:0040011), movement of cell/subcellular component (GO:0006928), and cell motility (GO:0048870) (Fig. 3a, Supporting Information Table 4), in agreement with the observance of numerous flagellin proteins significantly increased in our Rag samples. There was a significant enrichment of protein clusters associated with localization as well, in which the majority of the protein clusters associated with this GO term were transporters. Accordingly, molecular function (MF) GO terms for Rag samples showed a higher level of transporter activity (GO:0005215). The majority of the BP GO terms enriched in the WT samples are associated with RNA metabolic processes (including RNA processing (GO:0006396), RNA biosynthetic process (GO:0032774), mRNA metabolic process (GO:0016071), and ncRNA metabolic process (GO:0034660)) and translation (GO:0006412). Analysis of the MF GO terms changed in WT samples show a similar signature, with terms including, but not limited to, RNA polymerase activity (GO:0097747), DNA binding (GO:0003677), RNA binding (GO:0003723), and structural constituent of ribosome (GO:0003735). Overall, the Rag gut microbial proteome shows a vast increase in motility/flagellin proteins and transporter activity, whereas there is an enrichment in RNA and protein synthesis in the WT gut microbial proteome.

### 3.4 Protein Clusters Significantly Altered between Colitic and Healthy Mice

To distinguish potentially important proteins that are involved in intestinal inflammation and not due to the *Rag1*^-/-^ background (*i.e.,* baseline differences between the WT vs. Rag host proteins and/or gut microbiota), we focused our efforts on the identification of aberrant protein clusters in RT samples that were significantly altered between this treatment group and the combination of WT and Rag samples (hereafter called “controls”) (Fig. 2b, Fig. 3b, Supporting Information Tables 5, 6). 66 murine and 309 microbial protein clusters were significantly altered (FDR <0.05 and |log2FC| >1). Of the significantly altered murine protein clusters, 6 were increased in the control samples, and 54 were increased in the RT samples (Fig. 2b, Supporting Information Table 5). Interestingly, many of the murine proteins significantly increased in the RT samples are immune response-related genes, as would be expected with inflammation. Some examples include myeloperoxidase (MPO), granzyme A (GZMA), lactoferrin (LTF), and the pro-inflammatory proteins S100A8 and S100A9 [34]. Two of the most highly increased proteins are the two S100A proteins, which are abundantly found in neutrophils and other myeloid cells, and together form the damage-associated molecular pattern (DAMP) molecule calprotectin. This heterodimer has a wide variety of functions, including immune cell activation via binding and activation of toll like-receptor 4 (TLR4) and the receptor for advanced glycation end products (RAGE), calcium and zinc binding, and antimicrobial properties [34-36]. Elevated levels of calprotectin have been implicated in a variety of diseases such as cancer and IBD, and (along with MPO and LTF) has been used as a fecal marker of intestinal inflammation in IBD [37].

In addition, we observe an increase in many protease inhibitors, including several serpins (Serpina3k, Serpinc1, Serpina3f, Serping1, Serpina1a), MUG1, and PZP. This suggests an elevation of proteolytic activity that is usually associated with inflammation, and a host response attempting to suppress the elevated proteolytic activity. Furthermore, Serpina1 (α1-antitrypsin) has long been used as a fecal marker of intestinal inflammation in IBD [37,38].

It is well described that intestinal inflammation also results in epithelial responses. Various cytokines have been shown to increase intestinal epithelial cell production/secretion of antimicrobial peptides and mucus [39,40]. Accordingly, we observe an enrichment of several antimicrobial REG peptides (REG1, REG2, REG3A, REG3B) as well as MUC2 in the RT proteomic datasets. Together, this data shows that our metaproteomics methodology permits the identification and relative quantitation of many secreted host proteins known to be associated with intestinal inflammation.

Of the 309 significantly altered microbial protein clusters, 174 were increased in the control samples, and 135 were increased in the RT samples (Fig. 3b, Supporting Information Table 6). GO term analysis on the microbial proteins shows that the MF GO terms enriched in control samples include hydrolase activity acting on O-glycosyl bonds (GO:0004553), coenzyme binding (GO:0050662), and electron carrier activity (GO:0009055) (Supporting Information Table 7). In contrast, the MF GO terms in higher abundance in RT mice include RNA binding (GO:0003723), phosphoglycerate kinase activity (GO:0004618), and substrate-specific transporter activity (GO:0022892). Accordingly, we observe many phosphoglycerate kinases involved in glycolysis, as well as maltose transporters enriched in the RT samples.

Many of the significantly altered microbial proteins and GO terms enriched in the RT samples overlapped with functional signatures identified in either the Rag or WT samples. Therefore, to further interrogate these similarities and differentiate the functions unique to RT among the three cohorts, we performed additional comparisons between RT vs. WT (Fig. 2c, Fig. 3c, Supporting Information Tables 8-10) and RT vs. Rag (Fig. 2d, Fig. 3d, Supporting Information Tables 11-13). These analyses uncovered protein functionalities unique with respect to the RT samples, including the MF GO terms phosphoglycerate kinase (GO:0004618) and substrate-specific transporter activity (GO:0022892) as mentioned above, along with the BP GO terms monosaccharide metabolic process (GO:0005996) and single-organism localization (GO:1902578), among others. Notwithstanding, the RT mice shared enriched GO term functions with the Rag mice that were diminished and/or missing in WT (including locomotion (GO:0040011), movement of cell/subcellular component (GO:0006928), and cell motility (GO:0048870)), as well as numerous microbial protein functionalities matching the WT mice that were absent and/or reduced in the Rag group (namely those involved in transcription and translation).

### 3.5 Taxonomic Comparison with 16S rDNA Sequencing and Metaproteomic Mapping

We performed 16S rDNA sequencing on the same samples employed for proteomic analysis (Fig. 4a, Supporting Information Table 14). In agreement with the field, we observed that *Firmicutes* and *Bacteroidetes* comprised the majority of the commensal gut microbiota in the mice. Furthermore, we observed an altered ratio of these phyla in the RT mice, with an increase in *Proteobacteria* and *Verrucomicrobia,* which has been described in other mouse models of colitis as well as in IBD patients [3,12,41,42]. Importantly, these sequencing results correlate exceedingly well with the microbial composition analyses generated from our LC-MS/MS proteomics data at the phylum level (Fig. 4b). Importantly, the measurable increase in *Proteobacteria* and *Verrucomicrobia* in the RT samples was detected by both sequencing and proteomics methods. These results strongly support that the assessment of bacterial composition with proteomic data correlates well with the standard in the field of 16S sequencing.

**Fig. 4.**
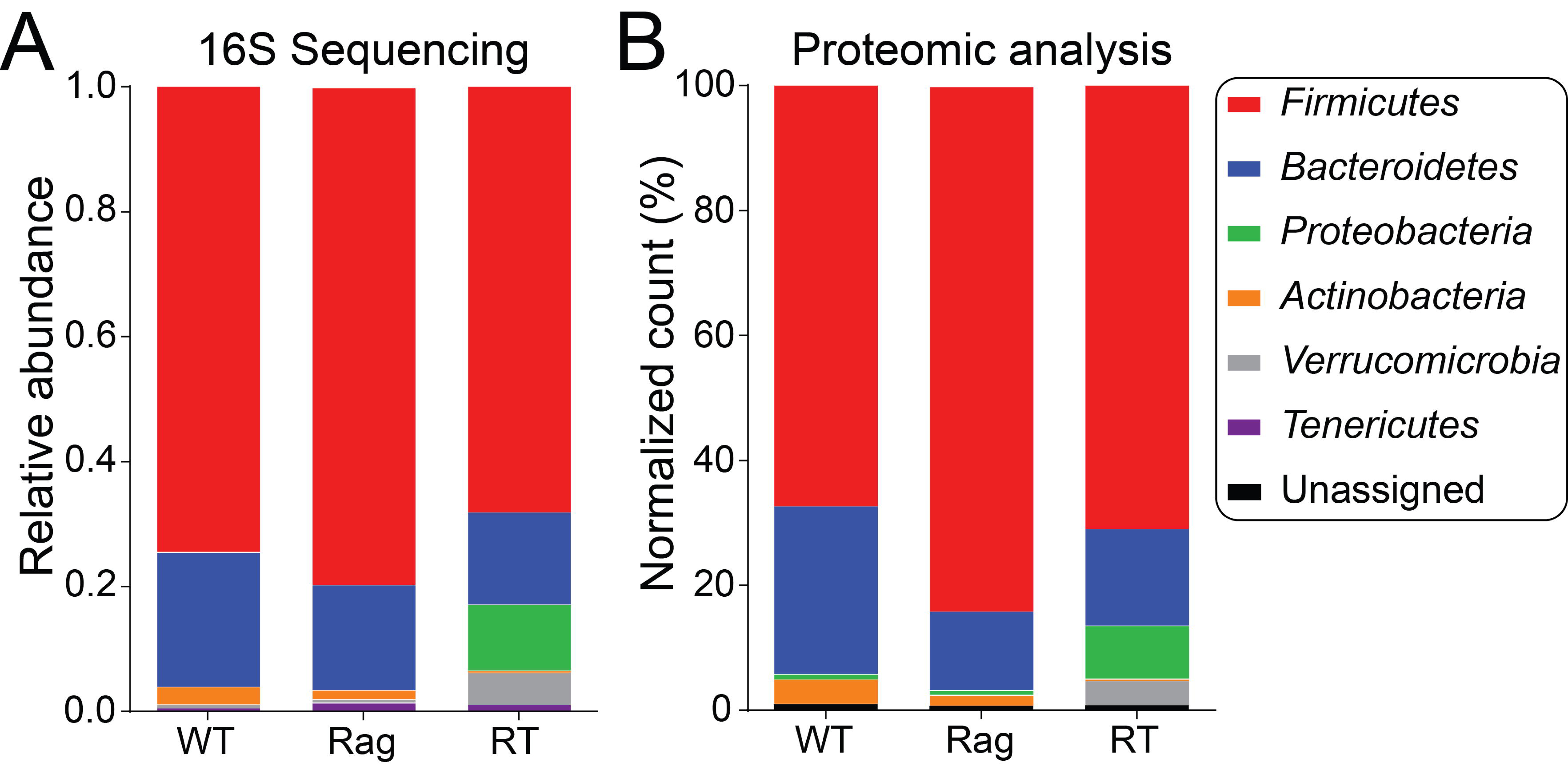
Taxonomic analyses of colonic bacteria. (A) Relative abundance of 16S rDNA sequence assignments of colonic bacterial samples at the phylum level. (B) Averaged peptide spectral counts attributable to a single phylum obtained from LC-MS/MS-based metaproteomics.

## 4 Discussion

For our T cell transfer colitis studies, we utilized 2 control groups to compare with the RT mice and help identify proteins important in colitis: 1) isogenic *Rag*^-/-^ mice that did not receive T cells and 2) C57BL6/J WT mice that have a commensal gut microbiota not altered by the lack of adaptive immune cells. Both PCA analysis and hierarchical clustering demonstrated distinct proteomic differences across these two control groups (Fig. 1c, Supporting Information Table 2) and analysis at the peptide level shows ∽50% overlap between Rag and WT samples (Fig. 1d). Among the most striking differences between the two control groups was the enrichment in motility/flagellar proteins in the Rag gut microbial proteome, compared to a robust enrichment in transcription- and translation-related proteins in the WT gut microbial proteome (Supporting Information Table 3). The majority of the significantly altered proteins that could be traced back to a specific bacterial phylum were mapped to *Firmicutes* and *Bacteroidetes,* similar to what we observed for the total taxonomic make-up by 16S sequencing and proteomic analyses (Fig. 4). These results show that while there are moderate changes in the microbial composition between the WT and immunodeficient Rag mice, there are tremendous alterations in microbial proteins and microbial protein functionalities represented in these mice, even in the absence of inflammation.

Similar to the striking locomotion/motility protein signature enriched in the Rag mice compared to WT, the RT mice also showed an abundance of flagellar proteins compared to WT mice (Supporting Information Tables 9, 10). Previous studies using *Tlr5*^-/-^ mice demonstrated that the absence of TLR5 resulted in the lack of flagellin-specific immunoglobulin, which allowed for the over-expression of flagellin by commensals [43]. Conversely, the presence of the anti-flagellin antibodies inhibited microbial motility, leading to the downregulation of flagellar proteins by the bacteria. A similar mechanism may be occurring in the Rag and RT mice in our studies, with the lack of B cells (and therefore immunoglobulins) leading to an increase of the normally low flagellin levels. Analysis of the murine proteins identified by our proteomic analyses show a B cell-deficient signature as expected, with a significant and notable absence of immunoglobulin proteins in the Rag and RT samples compared to WT. Interestingly, bacterial flagellins have been associated with pathogenicity, where pathogenic bacteria produce flagella to promote colonization and invasion of the mucosa [44]. The increased expression of bacterial flagellins in these mice may be another indication of a change in the commensal microbial population towards a more dysbiotic state. Furthermore, an increase in anti-flagellar antibodies have been associated with IBD, specifically CD [45], demonstrating a host response to this immune-reactive protein. Future studies utilizing a co-transfer of B cells in this T cell transfer colitis model can help distinguish the contributions of intestinal inflammation vs. genetic background in the expansion of bacterial flagellins in these mice.

Analysis of microbial protein-associated GO terms also revealed prominent similarities between the RT and WT samples, particularly with respect to GO terms pertaining to RNA and protein synthesis (Supporting Information Tables 4, 7, 13). We posit that the GO terms shared among the RT and WT mice are contributable to bacterial growth; however, the parental bacteria synthesizing the proteins are vastly different between the two groups. Both metaproteomics and 16S sequencing revealed that the inflammatory state of RT mice significantly alters the microbial phylogenetic composition, with a significantly increased footprint of *Proteobacteria* in the RT samples (Fig. 4). This phylum is known to expand in other intestinal inflammation models [3,12]. Therefore, the overabundance of microbial proteins involved in bacterial replication and protein synthesis likely represent the ability of *Proteobacteria* to thrive in the inflammatory state of RT mice, as previously shown [46]. Conversely, these same protein functionalities that dominate the microbial proteome in WT mice during homeostasis are attributable to the commensal organisms belonging to the *Firmicutes* and *Bacteroidetes* phyla (Fig. 3). In support of this hypothesis, we find that the majority of the bacterial proteins associated with these GO terms in the RT samples can be mapped back to *Proteobacteria,* while the WT sample proteins associated with these GO terms can be mapped to *Firmicutes* and *Bacteroidetes* (Supporting Information Tables 3, 6, 12). We do not observe an increase in transcriptional/translational proteins from the Rag microbiota relative to WT and RT mice, and this finding may be due to the significant differences in flagellar proteins that supersede the identification of those proteins associated with growth and division.

Analysis of the multiple comparisons identified protein functionalities unique to the RT group as well. As would be expected, many murine proteins associated with inflammation were significantly elevated in the RT group compared to either of the control groups, including both subunits of calprotectin, several protease inhibitors, and a variety of antimicrobial peptides (Supporting Information Tables 5, 8, 11). In addition, microbial protein functionalities were found specifically enriched in the RT group as well, including the MF GO term substrate-specific transporter activity (Supporting Information Tables 7, 10, 13). One class of proteins that corresponded to this enriched GO term included the maltose transporters (Supporting Information Tables 6, 9, 12). Studies elucidating the maltose transport system using *E. coli* have identified several components involved in the transport and utilization of maltose and maltodextrins, which is ultimately broken down to glucose by several enzymes [47,48]. Furthermore, glucose starvation leads to elevated expression of the maltose system genes, whereas bacterial cultures grown in high glucose concentrations leads to the block in expression of the *mal* genes [47]. The enrichment of maltose transporters and others in the RT associated bacteria may be an indicator of an increased need for or utilization of maltose and/or glucose in the inflamed state.

Altogether, these results demonstrate the important contributions of host genetics, the gut microbiota, and inflammation to the altered intestinal proteome observed in the RT mice, and support the combined use of 16S sequencing and metaproteomics methods incorporating an independent ComPIL protein database to microbiome studies [20]. These data provide a foundation for studying the alterations in intestinal luminal protein functionalities in a robust and well-characterized mouse model of colitis, and set the stage for future studies to further explore the functional mechanisms of some of these altered protein functionalities.

*We thank Ms. Ana Wang for assistance with mass spectrometry sample preparation protocols; Dr. Oktay Kirak for assistance with T cell transfer protocols; Dr. Robin Park for maintenance of Blazmass; and Dr. James Moresco, Ms. Jolene Diedrich, Ms. Barbara Orelo, and Prof. John Yates for technical assistance with mass spectrometry instrumentation; Mr. Mike Mayers for assistance with scripts for bioinformatic analyses. This work was supported by The Scripps Research Institute and the US National Institutes of Health 1R21CA181027 to D.W.W., U54GM114833 to A.I.S., and NIH Training Grant T32AI007244 to C.M.*

*The authors declare that they have no competing interests.*

## References

[1] Cadwell, K., Patel, K.K., Maloney, N.S., Liu, T.C., Ng, A., Virus-plus-susceptibility gene interaction determines Crohn’s disease gene Atg16L1 phenotypes in intestine. Cell 2010, 141, 1135–1145.

[2] Rogler, G., Interaction between susceptibility and environment: examples from the digestive tract. Dig. Diseases 2011, 29, 136–143.

[3] Elson, C.O., Cong, Y., Host-microbiota interactions in inflammatory bowel disease. Gut Microbes 2012, 3, 332–344.

[4] Leone, V., Chang, E.B., Devkota, S., Diet, microbes, and host genetics: the perfect storm in inflammatory bowel diseases. J. Gastroenterol. 2013, 48, 315–321.

[5] Graham, D.B., Xavier, R.J., From genetics of inflammatory bowel disease towards mechanistic insights. Trends Immunol. 2013, 34, 371–378.

[6] Xavier, R.J., Podolsky, D.K., Unravelling the pathogenesis of inflammatory bowel disease. Nature 2007, 448, 427–434.

[7] Kang, S.S., Bloom, S.M., Norian, L.A., Geske, M.J., et al, An antibiotic-responsive mouse model of fulminant ulcerative colitis. PLoS Med. 2008, 5, e41.

[8] Elson, C.O., Cong, Y., McCracken, V.J., Dimmitt, R.A., et al, Experimental models of inflammatory bowel disease reveal innate, adaptive, and regulatory mechanisms of host dialogue with the microbiota. Immunol. Rev. 2005, 206, 260–276.

[9] Sartor, R.B., Therapeutic correction of bacterial dysbiosis discovered by molecular techniques. Proc. Nat. Acad. Sci. USA 2008, 105, 16413–16414.

[10] Dalal, S.R., Chang, E.B., The microbial basis of inflammatory bowel diseases. J. Clin. Invest. 2014, 124, 4190–4196.

[11] Huttenhower, C., Kostic, A.D., Xavier, R.J., Inflammatory bowel disease as a model for translating the microbiome. Immunity 2014, 40, 843–854.

[12] Frank, D.N., St Amand, A.L., Feldman, R.A., Boedeker, E.C., et al, Molecular-phylogenetic characterization of microbial community imbalances in human inflammatory bowel diseases. Proc. Nat. Acad. Sci. USA 2007, 104, 13780–13785.

[13] Human Microbiome Project Consortium, Structure, function and diversity of the healthy human microbiome. Nature 2012, 486, 207–214.

[14] Lozupone, C.A., Stombaugh, J.I., Gordon, J.I., Jansson, J.K., Knight, R., Diversity, stability and resilience of the human gut microbiota. Nature 2012, 489, 220–230.

[15] Kolmeder, C.A.C., de Been, M.M., Nikkilä, J.J., Ritamo, I.I., et al, Comparative metaproteomics and diversity analysis of human intestinal microbiota testifies for its temporal stability and expression of core functions. PLoS ONE 2012, 7, e29913–e29913.

[16] Li, X.X., Leblanc, J.J., Truong, A.A., Vuthoori, R.R., et al, A metaproteomic approach to study human-microbial ecosystems at the mucosal luminal interface. PLoS ONE 2011, 6, e26542–e26542.

[17] Verberkmoes, N.C., Russell, A.L., Shah, M., Godzik, A., et al, Shotgun metaproteomics of the human distal gut microbiota. ISME J. 2009, 3, 179–189.

[18] Erickson, A.R., Cantarel, B.L., Lamendella, R., Darzi, Y., et al, Integrated metagenomics/metaproteomics reveals human host-microbiota signatures of Crohn’s disease. PLoS ONE 2012, 7, e49138.

[19] Wolters, D.A.D., Washburn, M.P.M., Yates, J.R.J., An automated multidimensional protein identification technology for shotgun proteomics. Anal. Chem. 2001, 73, 5683–5690.

[20] Chatterjee, S., Stupp, G.S., Park, S.K.R., Ducom, J.-C., et al, A comprehensive and scalable database search system for metaproteomics. BMC Genomics 2016, 17, 642.

[21] Powrie, F., Leach, M.W., Mauze, S., Caddie, L.B., Coffman, R.L., Phenotypically distinct subsets of CD4+ T cells induce or protect from chronic intestinal inflammation in C. B-17 scid mice. Int. Immun. 1993, 5, 1461–1471.

[22] Ostanin, D.V., Bao, J., Koboziev, I., Gray, L., et al, T cell transfer model of chronic colitis: concepts, considerations, and tricks of the trade. Am. J. Physiol. Gastrointest. Liver Physiol. 2009, 296, G135–G146.

[23] Mayers, M.D., Moon, C., Stupp, G.S., Su, A.I., Wolan, D.W., Quantitative metaproteomics and activity-based probe enrichment reveals significant alterations in protein expression from a mouse model of inflammatory bowel disease. J. Proteome Res. 2017, 16, 1014–1026.

[24] Wu, J., Zhu, J., Yin, H., Liu, X., et al, Development of an integrated pipeline for profiling microbial proteins from mouse fecal samples by LC-MS/MS. J. Proteome Res. 2016, 15, 3635–3642.

[25] Elias, J.E., Gygi, S.P., Target-decoy search strategy for increased confidence in large-scale protein identifications by mass spectrometry. Nat. Methods 2007, 4, 207–214.

[26] Love, M.I., Huber, W., Anders, S., Moderated estimation of fold change and dispersion for RNA-seq data with DESeq2. Genome Biol. 2014, 15, 550.

[27] Subramanian, A., Tamayo, P., Mootha, V.K., Mukherjee, S., et al, Gene set enrichment analysis: a knowledge-based approach for interpreting genome-wide expression profiles. Proc. Nat. Acad. Sci. USA 2005, 102, 15545–15550.

[28] Caporaso, J.G., Kuczynski, J., Stombaugh, J., Bittinger, K., et al, QIIME allows analysis of high-throughput community sequencing data. Nat. Chem. Biol. 2010, 7, 335–336.

[29] Masella, A.P., Bartram, A.K., Truszkowski, J.M., Brown, D.G., Neufeld, J.D., PANDAseq: paired-end assembler for Illumina sequences. BMC Bioinformatics 2012, 13, 31.

[30] DeSantis, T.Z., Hugenholtz, P., Larsen, N., Rojas, M., et al, Greengenes, a chimera-checked 16S rRNA gene database and workbench compatible with ARB. Appl. Environ. Microbiol. 2006, 72, 5069–5072.

[31] Lozupone, C., Knight, R., UniFrac: a new phylogenetic method for comparing microbial communities. Appl. Environ. Microbiol. 2005, 71, 8228–8235.

[32] Vizcaíno, J.A., Csordas, A., Del-Toro, N., Dianes, J.A., et al, 2016 update of the PRIDE database and its related tools. Nucleic Acids Res. 2016, 44, 11033–11033.

[33] Langley, S.R., Mayr, M., Comparative analysis of statistical methods used for detecting differential expression in label-free mass spectrometry proteomics. J. Proteomics 2015, 129, 83–92.

[34] Vogl, T., Gharibyan, A.L., Morozova-Roche, L.A., Pro-inflammatory S100A8 and S100A9 proteins: self-assembly into multifunctional native and amyloid complexes. Int. J. Mol. Sci. 2012, 13, 2893–2917.

[35] Stríz, I., Trebichavský, I., Calprotectin - a pleiotropic molecule in acute and chronic inflammation. Physiol. Res. 2004, 53, 245–253.

[36] Srikrishna, G., S100A8 and S100A9: new insights into their roles in malignancy. J. Innate Immun. 2012, 4, 31–40.

[37] Foell, D., Wittkowski, H., Roth, J., Monitoring disease activity by stool analyses: from occult blood to molecular markers of intestinal inflammation and damage. Gut 2009, 58, 859–868.

[38] Fischbach, W., Becker, W., Mössner, J., Koch, W., Reiners, C., Faecal alpha-1-antitrypsin and excretion of ^111^indium granulocytes in assessment of disease activity in chronic inflammatory bowel diseasess. Gut 1987, 28, 386–393.

[39] Onyiah, J.C., Colgan, S.P., Cytokine responses and epithelial function in the intestinal mucosa. Cell. Mol. Life Sci. 2016, 73, 4203–4212.

[40] Mukherjee, S., Hooper, L.V., Antimicrobial defense of the intestine. Immunity 2015, 42, 28–39.

[41] Berry, D., Kuzyk, O., Rauch, I., Heider, S., et al, Intestinal microbiota signatures associated with inflammation history in mice experiencing recurring colitis. Front. Microbiol. 2015, 6, 1408.

[42] Berry, D., Schwab, C., Milinovich, G., Reichert, J., et al, Phylotype-level 16S rRNA analysis reveals new bacterial indicators of health state in acute murine colitis. ISME J. 2012, 6, 2091–2106.

[43] Cullender, T.C., Chassaing, B., Janzon, A., Kumar, K., Innate and adaptive immunity interact to quench microbiome flagellar motility in the gut. Cell Host Microbe 2013, 14, 571–581.

[44] Ramos, H.C., Rumbo, M., Sirard, J.-C., Bacterial flagellins: mediators of pathogenicity and host immune responses in mucosa. Trends Microbiol. 2004, 12, 509–517.

[45] Alexander, K.L., Targan, S.R., Elson, C.O., Microbiota activation and regulation of innate and adaptive immunity. Immunol. Rev. 2014, 260, 206–220.

[46] Winter, S.E., Winter, M.G., Xavier, M.N., Thiennimitr, P., et al, Host-derived nitrate boosts growth of *E. coli* in the inflamed gut. Science 2013, 339, 708–711.

[47] Boos, W., Shuman, H., Maltose/maltodextrin system of *Escherichia coli*: transport, metabolism, and regulation. Microbiol. Mol. Biol. Rev. 1998, 62, 204–229.

[48] Nikaido, H., Maltose transport system of *Escherichia coli*: An ABC type transporter. FEBS Lett. 1994, 346, 55–58.

